## Appendix 2 - Taxon-specific richness maps and comments for "No one-size-fits-all solution to clean GBIF"

### Appendix 2 - No one-size-fits-all solution to clean GBIF

Alexander Zizka^1,2^, Fernanda Antunes Carvalho^3^, Alice Calvente^4^, Mabel Rocio Baez-Lizarazo^5^, Andressa Cabral^6^, Jéssica Fernanda Ramos Coelho^4^, Matheus Colli-Silva^6^, Mariana Ramos Fantinati^4^, Moabe Ferreira Fernandes^7^, Thais Ferreira-Araújo^4^, Fernanda Gondim Lambert Moreira^4^, Nathália Michelly da Cunha Santos^4^, Tiago Andrade Borges Santos^7^, Renata Clicia dos Santos‐Costa^4^, Filipe Cabreirinha Serrano^8^, Ana Paula Alves da Silva^4^, Arthur de Souza Soares^4^, Paolla Gabryelle Cavalcante de Souza^4^, Eduardo Calisto Tomaz^4^, Valéria Fonseca Vale^4^, Tiago Luiz Vieira^7^, Alexandre Antonelli^9,10,11^

1. sDiv, German Center for Integrative Biodiversity Research Halle-Jena-Leipzig (iDiv), Leipzig, Germany
2. Naturalis Biodiversity Center, Leiden, The Netherlands
3. Departamento de Genética, Ecologia e Evolução, Universidade Federal de Minas Gerais, Belo Horizonte, Brazil
4. Departamento de Botânica e Zoologia, Universidade Federal do Rio Grande do Norte, Natal, Brazil
5. Universidade Federal do Rio Grande do Sul, Porto Alegre, Brazil
6. Departamento de Botânica, Universidade de São Paulo, São Paulo, Brazil
7. Departamento de Ciências Biológicas, Universidade Estadual de Feira de Santana, Feira de Santana, Brazil
8. Departamento de Ecologia, Universidade de São Paulo, São Paulo, Brazil
9. Gothenburg Global Biodiversity Centre, University of Gothenburg, Gothenburg, Sweden
10. Department for Biological and Environmental Sciences, University of Gothenburg, Gothenburg, Sweden
11. Royal Botanic Gardens Kew, Richmond, United Kingdom

### Arhynchobatidae Fowler, 1934

The Arhynchobatidae family comprises 13 genera with approximately 104 species of softnose skates occurring in all oceans (Last *et al*., 2016). The species richness map of filtered data is more informative in comparison to the species richness map of raw data, albeit neither of them is likely to correctly represent the true richness of arhynchobatids in the Neotropics. According to Weigmann’s checklist (2016), it is clear that both raw and filtered assessments underestimate arhynchobatid richness. For example, the south-western Atlantic and south-eastern Pacific regions together hold 46 arhynchobatid species (Weigmann, 2016), yet less than 20 species are present in both raw and filtered richness maps for the same areas. This result is unexpected given that the number of flagged records removed from the analysis was relatively small.

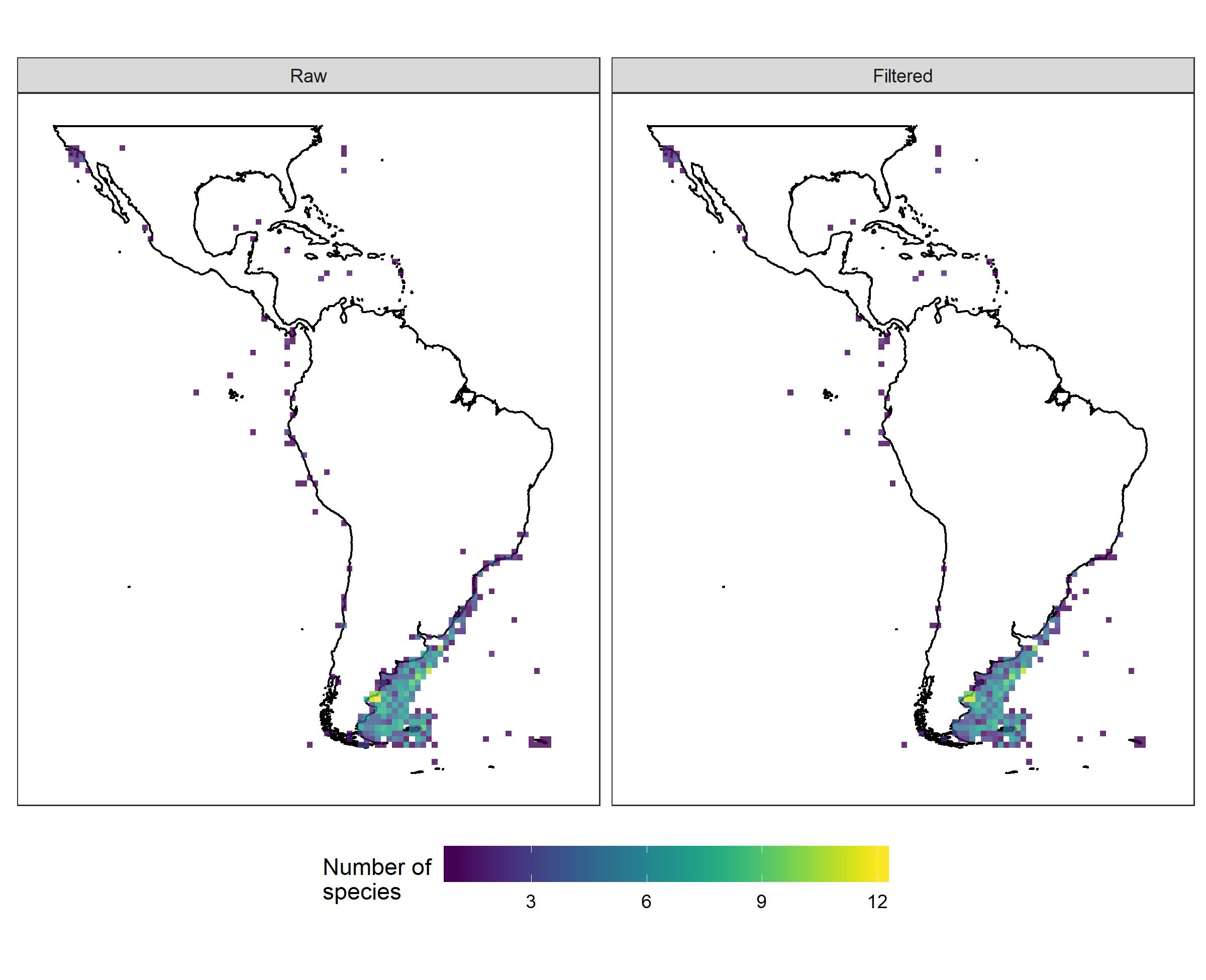

### *Conchocarpus* J.C.Mikan

The raw richness map for *Conchocarpus* – a neotropical genus of flowering plants with c. 50 species typical to the Amazon and Atlantic rainforests – shows a particular concentration of species in central Brazil. The number of records decreases by 43% when comparing the raw with the filtered dataset; conversely, the number of taxa does not fall significantly, decreasing in only one species between the two datasets. The flagged records in central Brazil represent mostly country centroids, and hence refer to imprecise occurrences. The raw map shows several duplicated records. Therefore, the filtered database is more accurate and closely represents the known distribution of the group (considering works such as Kallunki & Pirani, 1998; Colli-Silva & Pirani, 2019).

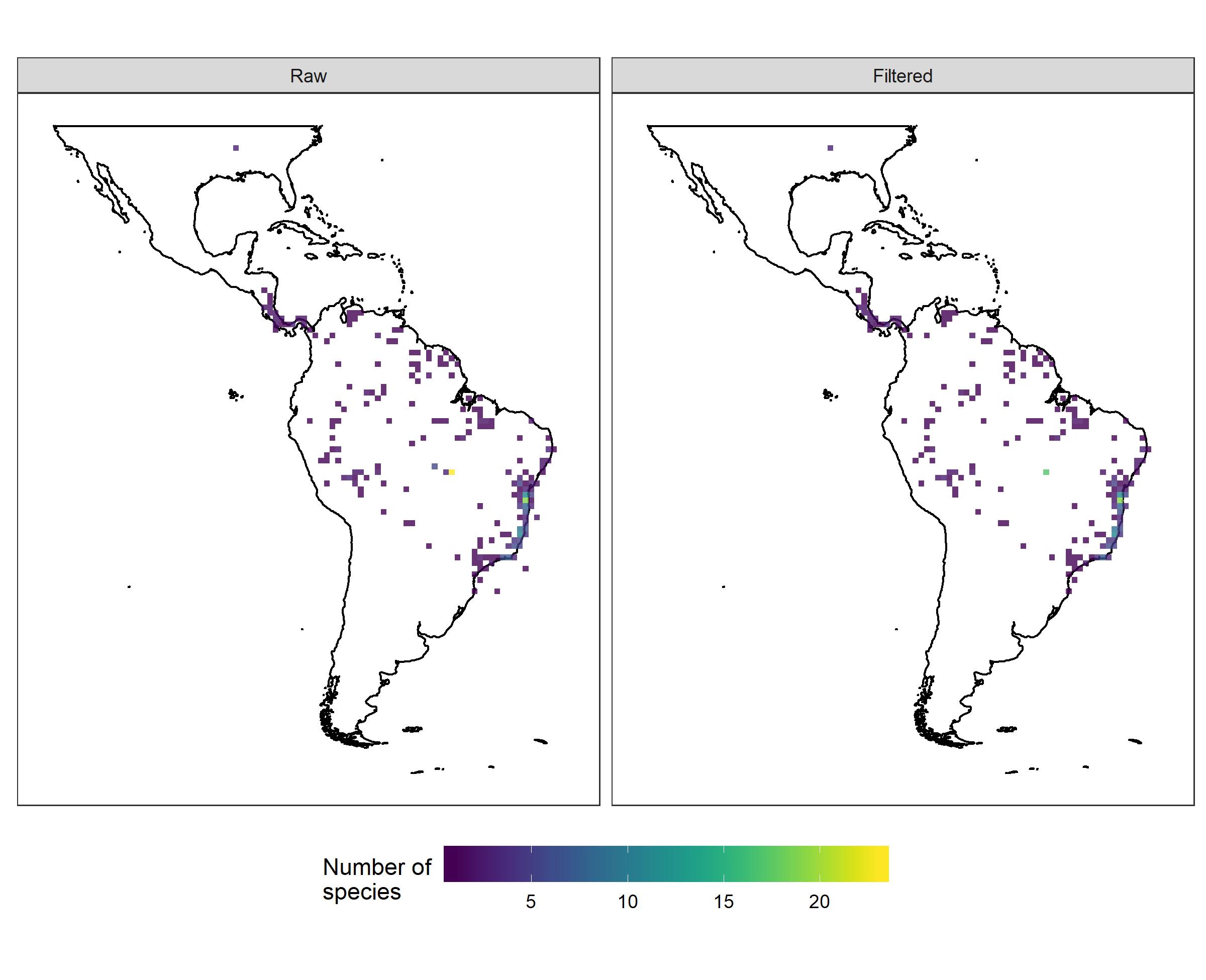

### Diogenidae Ortmann, 1892

The richness maps using raw and filtered data are similar at a large scale (gamma diversity). Both maps show higher number of species along the North and Central American rocky coasts (California, Gulf of Mexico, Florida, Panama and Caribbean) corresponding to the sea ecoregions “Tropical Eastern Pacific” and “Tropical Atlantic” (Spalding *et al*. 2007). Although Diogenidae are widespread in the pantropical oceans, there is an information gap in relation to hermit crabs in South America, and thus the number of records along these regions may be underestimated. (Vale *et al*. 2017). Historically, the few records in the north and northeast of Brazil may be a consequence of the scarcity of studies on the local aquatic biodiversity, especially for invertebrate groups.

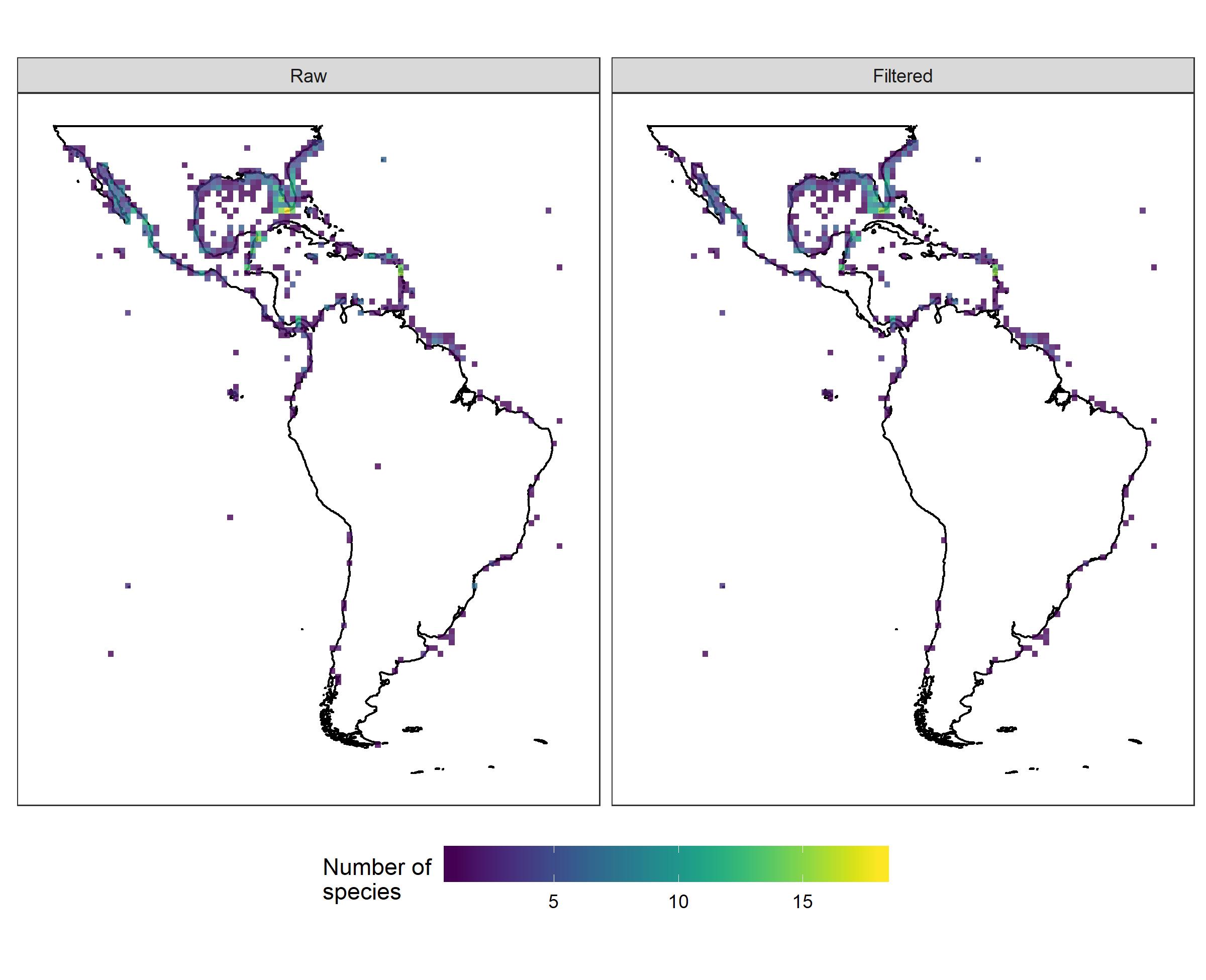

### Dipsadidae Bonaparte 1838

Dipsadidae is the richest snake clade in the New World, ranging from southern North America to the Patagonian steppes (Cadle & Greene, 1993). It comprises more than 700 species (Grazziotin et al., 2012) varying greatly in body size, diet and other natural history aspects (Cadle & Greene, 1993), with one genus (*Thermophis*) also occurring in Tibet (He et al., 2009). The high number of eliminated records was not expected and was mainly due to duplicate records. The overall richness patterns were similar between the raw and filtered datasets, with Central America, Ecuador and southeastern Brazil attaining the highest richness. Filtering eliminated several records in northern South America and erroneous sea records, as well as some potential collection-biased records in southeastern Brazil.

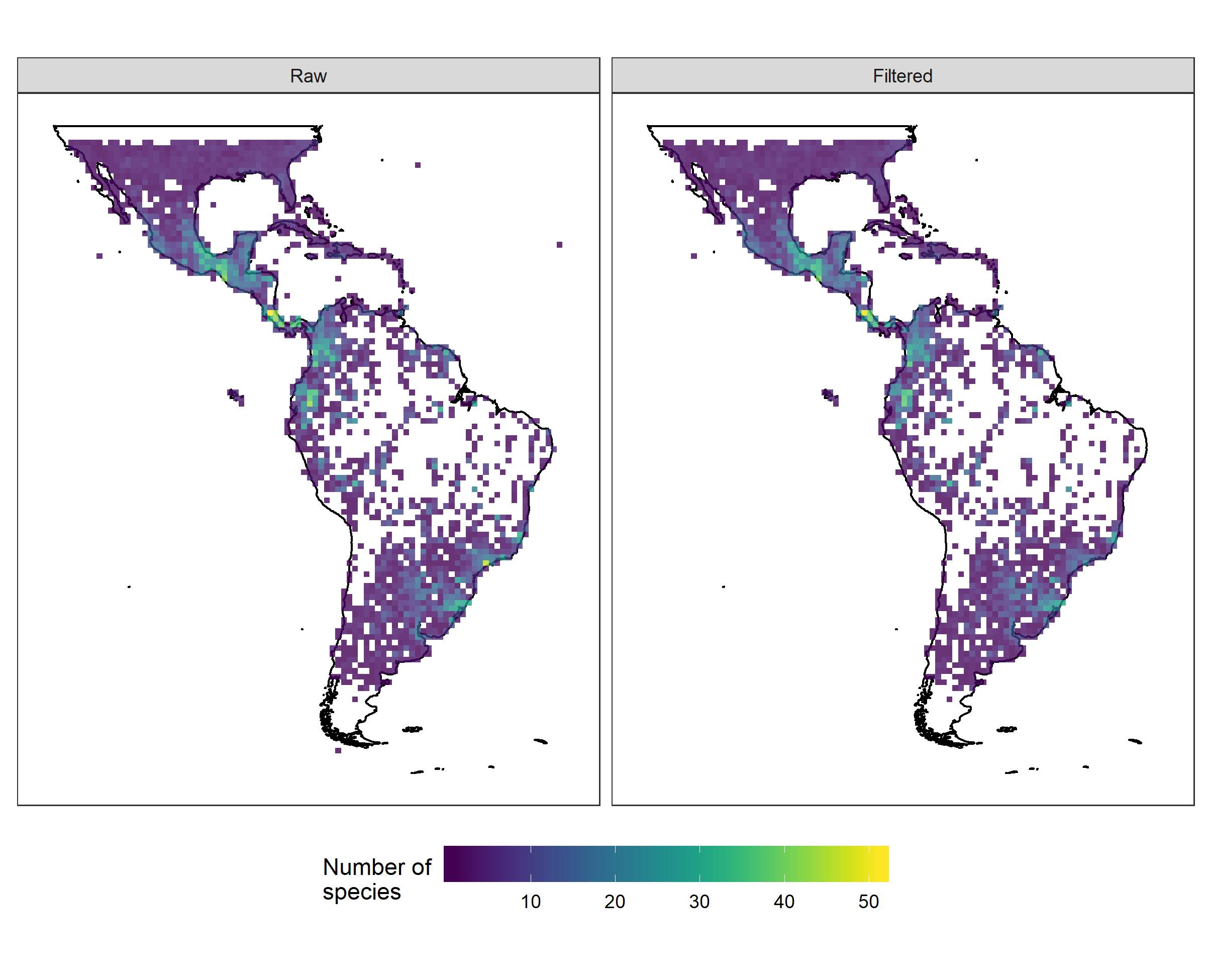

### Entomobryidae Tömösvary, 1882

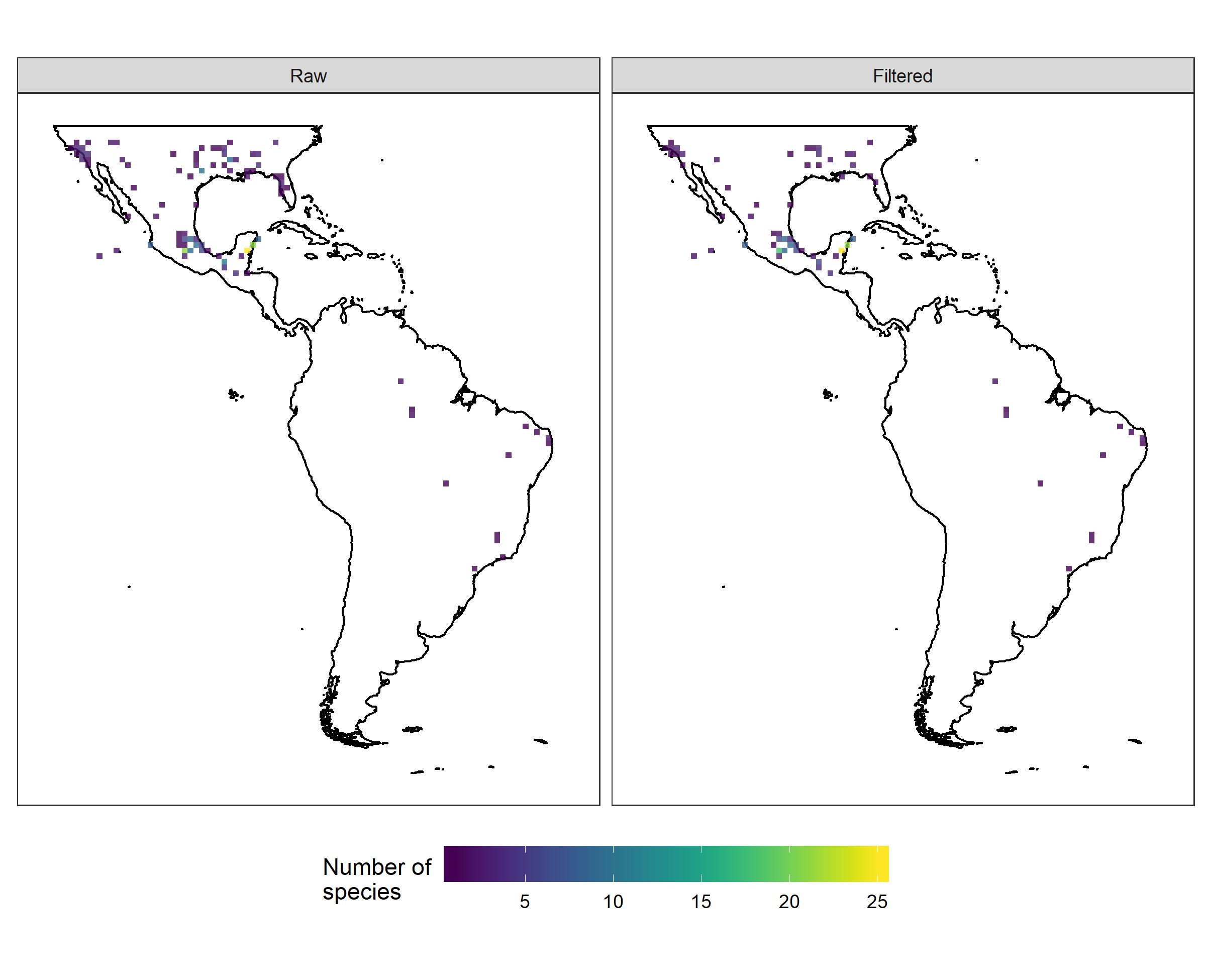

Entomobryidae is one of the richest families in Collembola, and this cosmopolitan taxon has approximately 174,120 records worldwide in the GBIF database. Of these, approximately 2,800 refer to the Neotropical region. The maps of richness, raw and filtered, obtained for this family are very similar, but they are not sufficiently demonstrative of the real richness and distribution of the taxon, possibly due to collection bias. It seems that the neotropical records for Entomobryidae are concentrated mainly where there are important Collembola research groups, such as in southern USA, Mexico (at the Universidad Nacional Autónoma de México) and Brazil (at the National Institute for Research in the Amazon and the Federal University of Rio Grande Do Norte). Most records flagged during the tests were duplicates.

### *Gaylussacia* Kunth

*Gaylussacia* (Ericaceae) currently comprises 54 species (Romão 2011), which occur in the subtropical portion of North America and tropical and subtropical South America; the genus is absent from Central America (Sleumer 1967; Floyd 2002; Romão 2011). The Brazilian Espinhaço Range, a Neotropical region located in Bahia and Minas Gerais states, is considered a center of diversity for *Gaylussacia* (Kinoshita-Gouvêa 1980). The filtered richness map was more accurate, and although it disregarded five taxa, it matched the current known distribution of the group (Romão 2011). As many species are endemic to Brazilian highlands and rocky fields (BFG 2018), the use of inaccurate geographic data could result in an increase in area of occurrence for *Gaylussacia* and consequently in an incorrect classification of the threat status. This possibly affected the distribution of five taxa not classified in the category "Endangered" in raw data evaluations, negatively affecting the conservation of these taxa and their habitats.

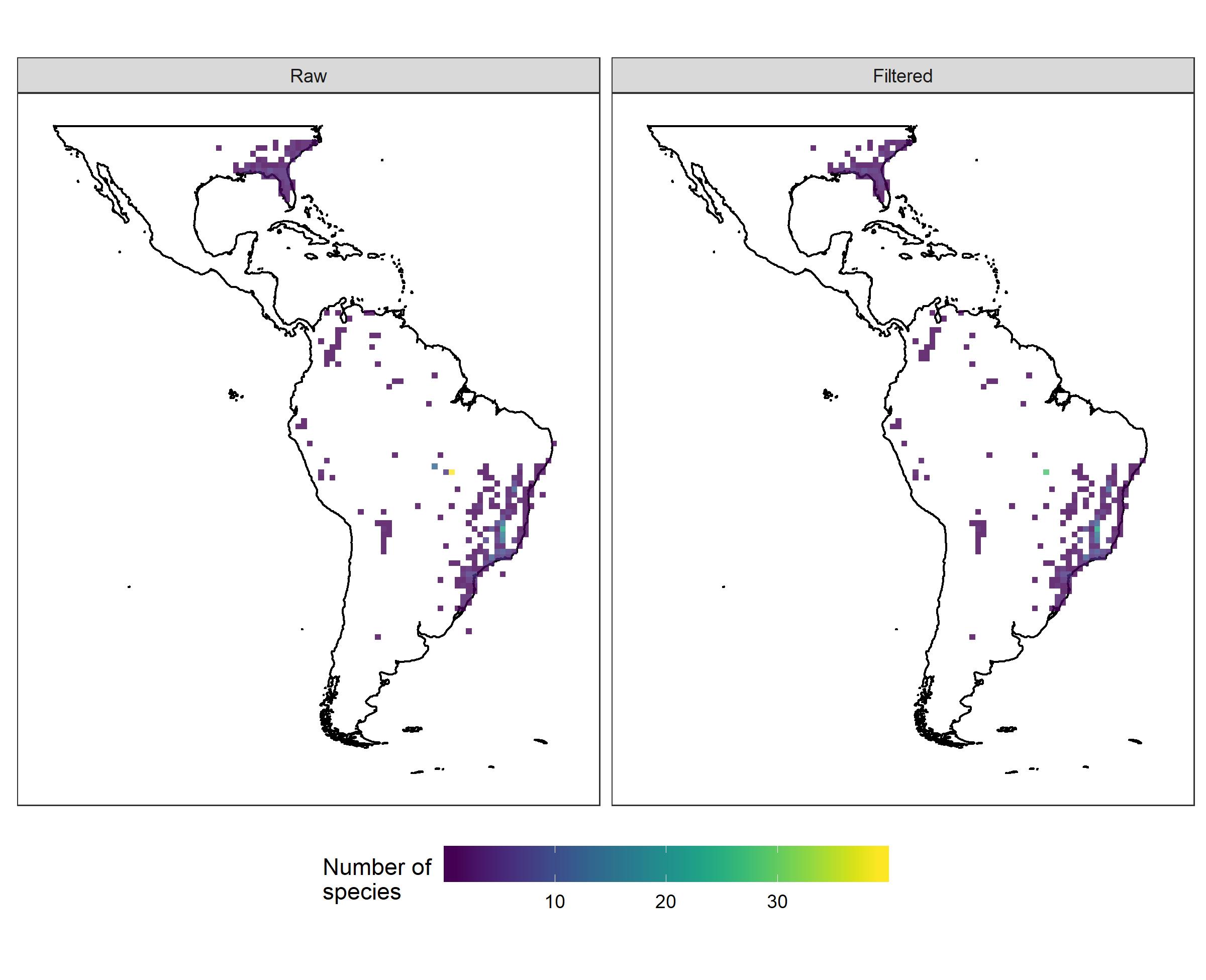

### *Harengula* Valenciennes 1847

*Harengula* is a genus of small coastal fish from the Clupeidae family (Clupeiformes), (Fricke et al. 2019; Whitehead 1985). This genus has four species and occurs in both the Pacific and Atlantic Oceans. The only species in the Pacific is *Harengula thrissina* (Jordan & Gilbert 1882), distributed from the Gulf of California (USA) to Peru (Whitehead 1985). In the Atlantic Ocean, there are *Harengula humeralis* (Cuvier 1829), *Harengula clupeola* (Cuvier 1829), and *Harengula jaguana* Poey 1865 (Fricke et al. 2019; Whitehead 1985). While *H. humeralis* only occurs from Florida (USA) to the Guianas, *H. clupeola* and *H. jaguana* co-occur from southern USA to southern Brazil (Whitehead 1985). Even after cleaning, there were suspicious records, such as inland occurrences. Filtered and raw richness maps were relatively similar, although the filtered richness map showed fewer inland occurrences records.

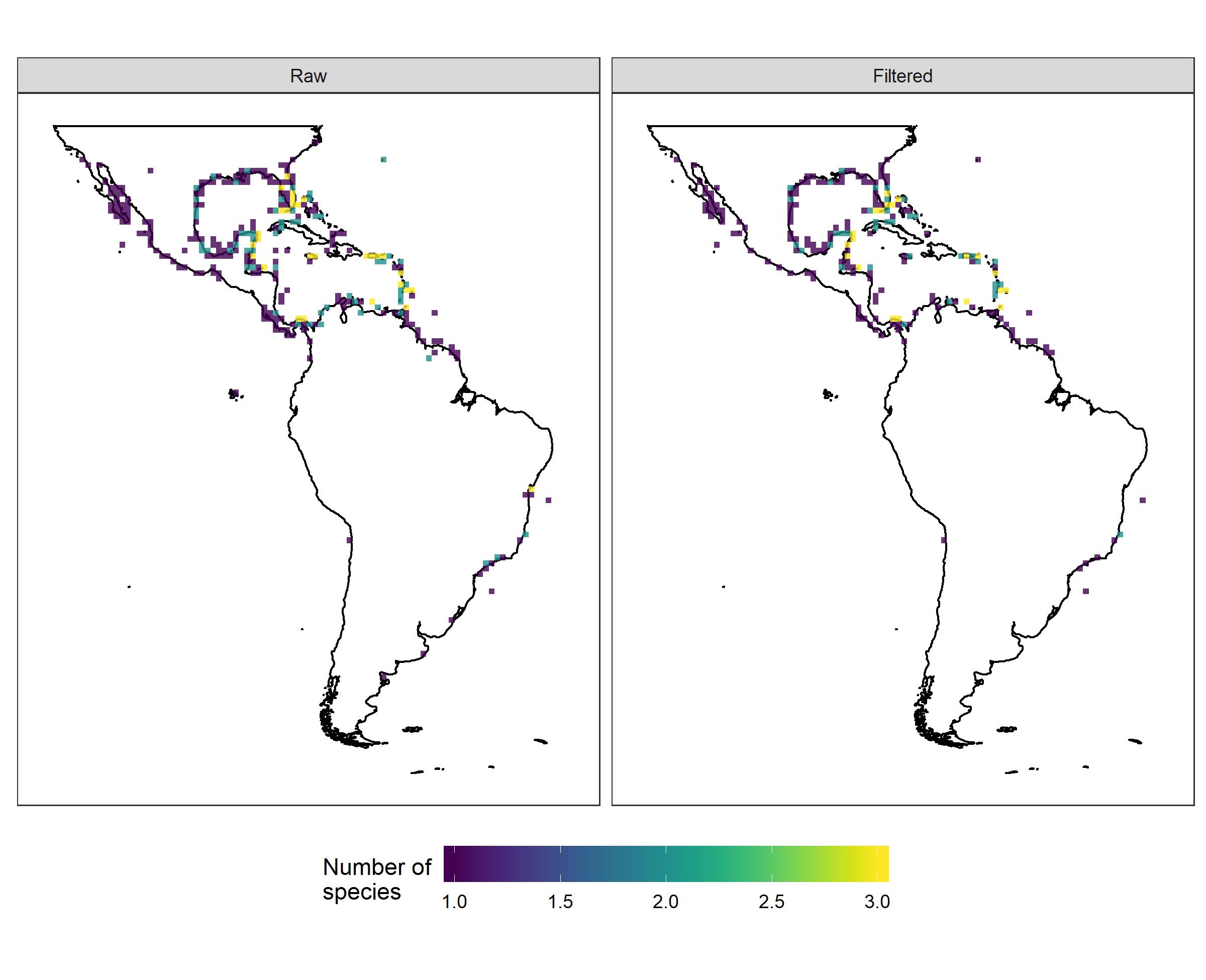

### *Harpalyce* Moç. & Sessé ex DC.

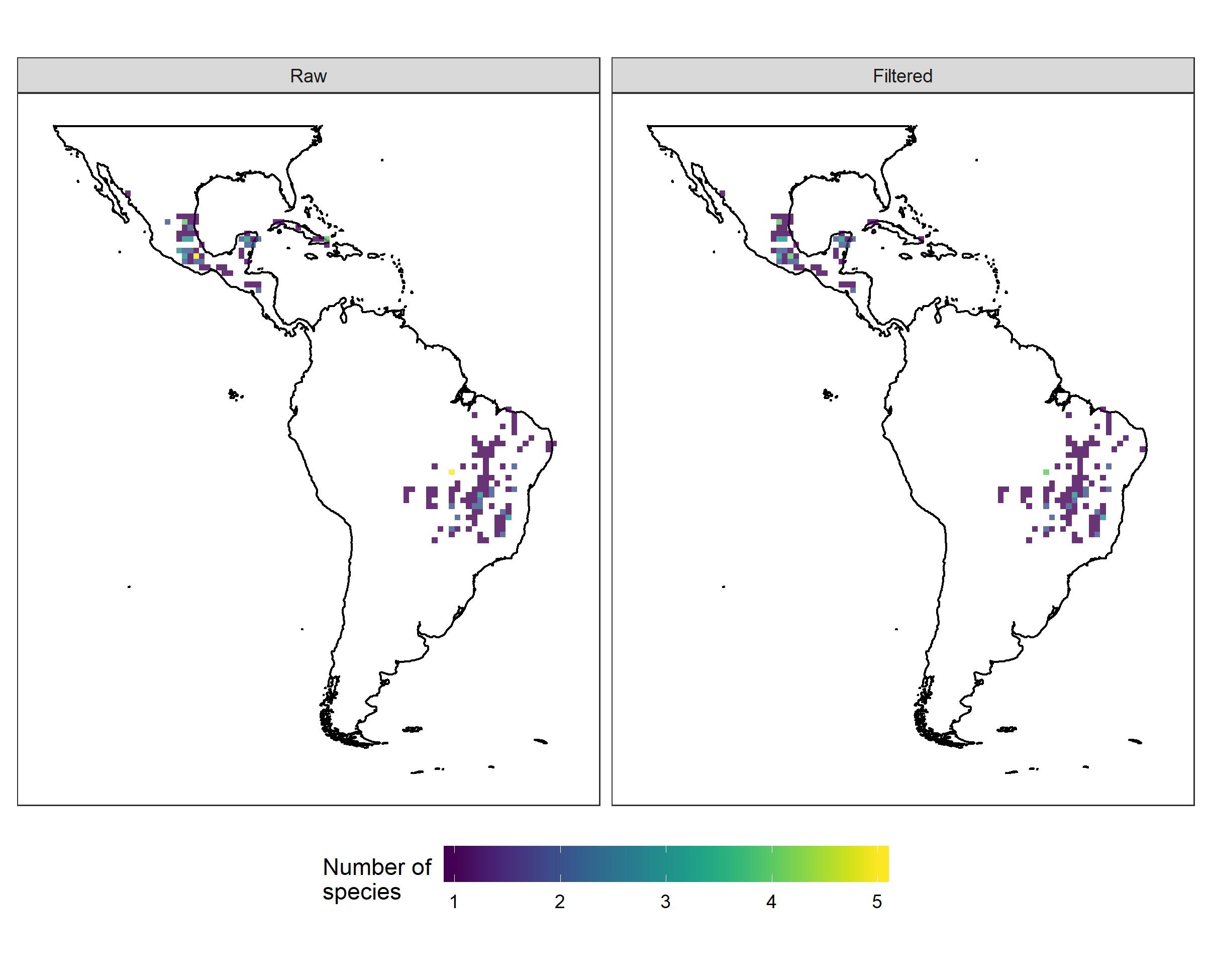

The genus *Harpalyce* (Fabaceae) comprises approximately 30 species, which are mostly associated with dry forests and savannas of the Neotropics (Arroyo 1976). Both maps recovered Mesoamerica and the Brazilian Central Plateau as areas of particularly high species richness, which is in line with the known distribution and the centres of endemism for *Harpalyce* (Arroyo 1976); these areas also harbour a significant number of recently described species (São-Mateus et al. 2016; 2018). The exceptional contribution of duplicated records (~50% of all flagged records) does not affect diversity patterns, and major differences between maps highlight distortions that may be caused by relatively few records. Country centroids, for example, generate two exceptionally rich grid cells in Brazil and Mexico.

### Iridaceae Juss.

The Neotropics are the second most important center of diversity for pantropical Iridaceae (Goldblatt & Manning, 2008). This family is represented in the Neotropical region by approximately 410 species within 25 genera (Goldblatt *et al.,* 2008). The flagged records in the family include mostly records in urban areas, duplicated records and those without species-level identification. This is reasonable and should be higher due to: registration of exotic ornamental species in datasets (of the genera *Aristea* Aiton, *Babiana* Ker Gawl.*, Chasmanthe* N.E.Brown, *Crocosmia* Planchon*, Dietes* Klatt*, Fosteria* Molseed*, Freesia* Klatt*, Gladiolus* L.*,* some *Iris* L.*, Moraea* Miller*, Romulea* Maratti and *Watsonia* Miller); spelling mistakes in taxon names (e.g. *Orthrosanthus chimboracensis* (Kunth) Baker, and *O. chimborasensis*); and the use of invalid species names (e.g. *Cardenanthus* R.C.Foster). The raw and filtered richness maps showed similar distribution patterns; however, when flags were eliminated, the most species-rich areas were better represented. Richness maps indicated that main diversity centers are located in Central Mexico, south and southeastern Brazil, followed by Andean South America (northern Andes to the Sub-Andes). These results concur with previous studies at tribal and generic levels (Trimezieae and Tigridieae, *Sisyrinchium* L.) (see Chauveau *et al.,* 2011; 2012; Souza-Chies *et al.,* 2012; Munguin-Lino *et al.,* 2016; Lovo *et al*., 2018).

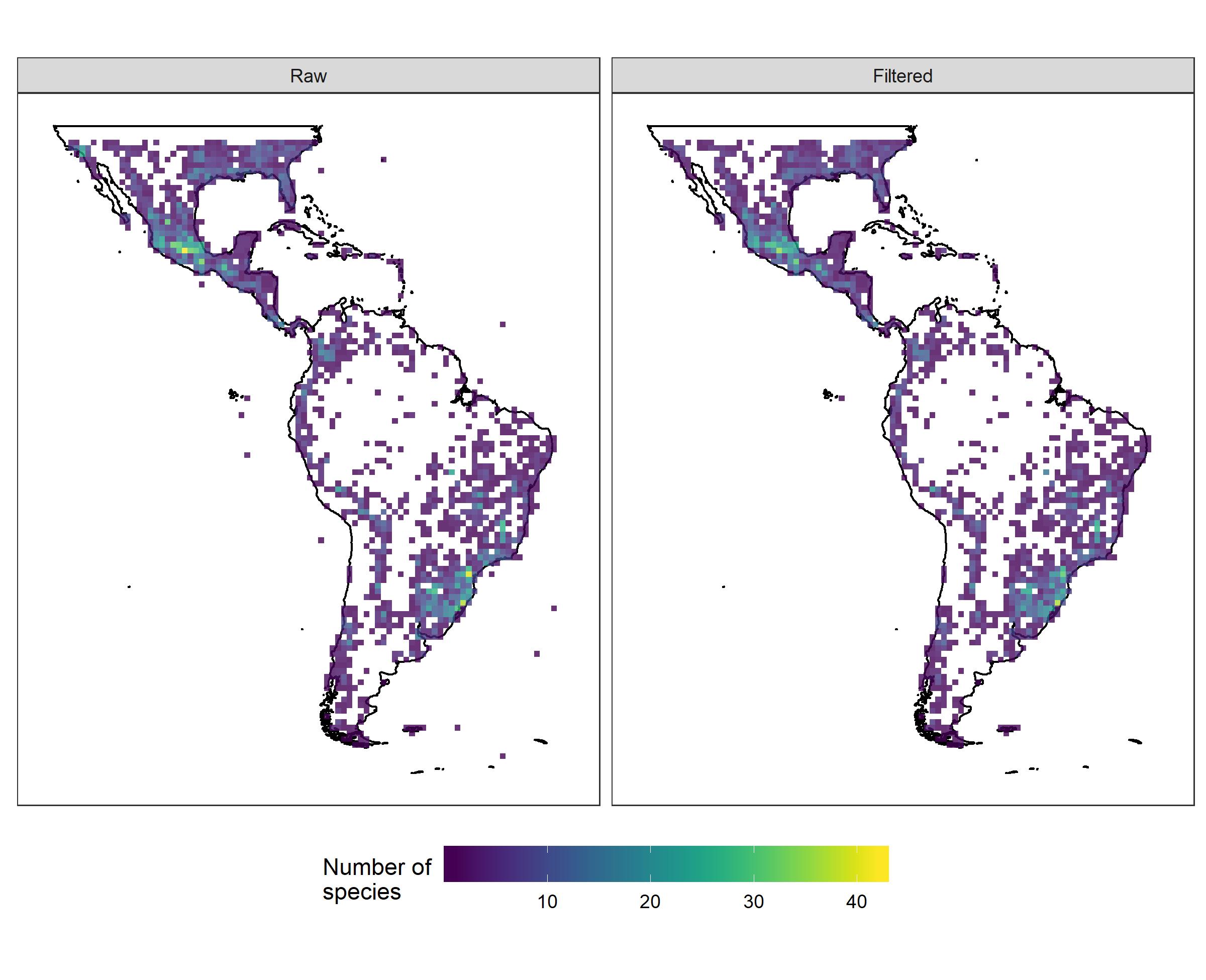

### *Lepismium* Pfeiff.

The epiphyte or rupicolous genus *Lepismium*, with six species, belongs to tribe Rhipsalideae (Cactaceae) and occurs exclusively in the Neotropical region (Nyffeler and Eggli et al. 2010). Richness maps based on raw and filtered data were quite similar and congruent with the previously known distribution (Hunt et al. 2006). *Lepismium* occurs along South America, with the highest number of species in Argentina followed by Bolivia and Brazil.

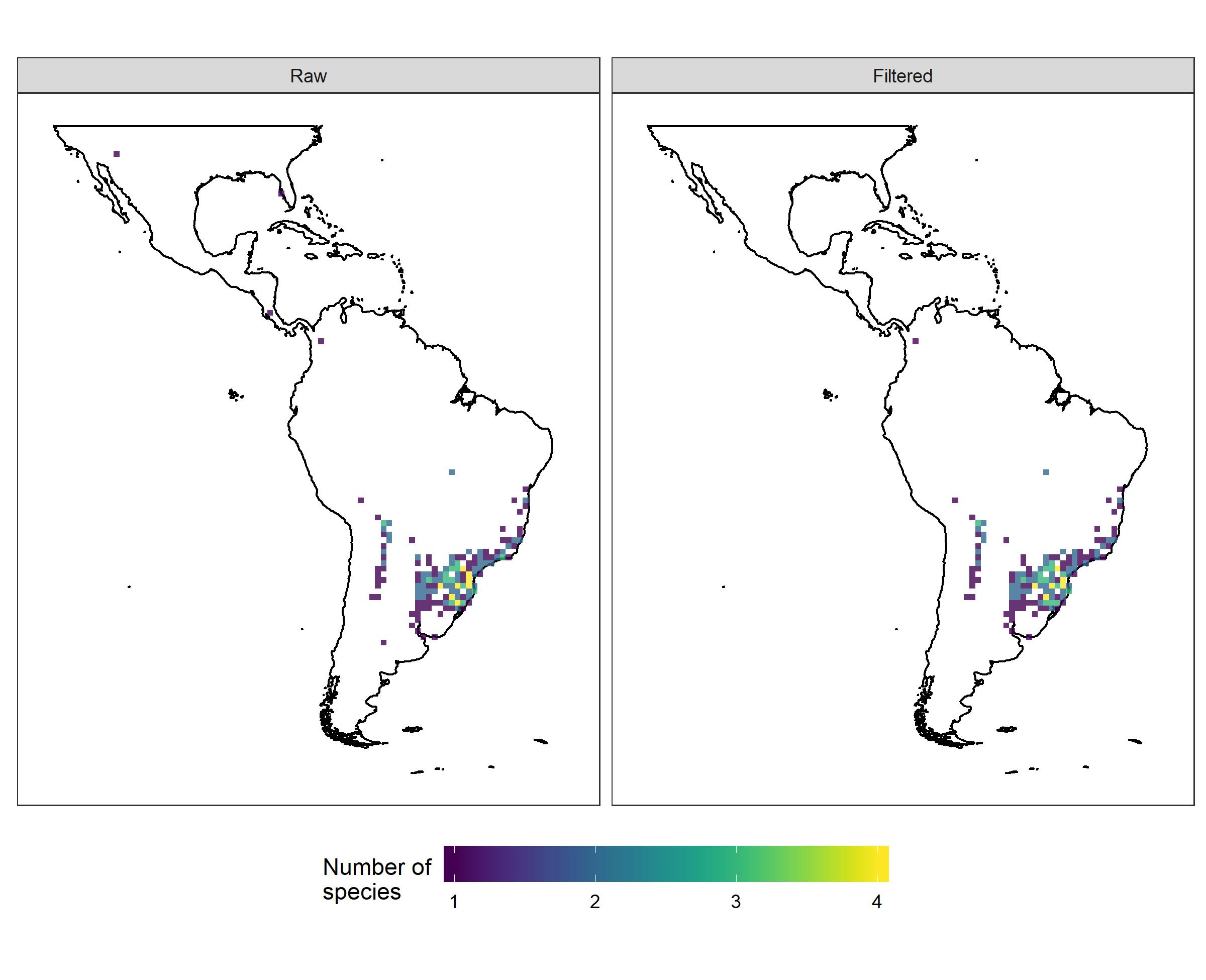

### Neanuridae Borner, 1901 *sensu* Yosii, 1956

Springtails (Collembola) are terrestrial microarthropods (0.12 to 17 mm in length), which live on wet soils and potentially represent a large part of neotropical diversity not yet deeply investigated. There are approximately 8,800 described species, occurring across all continents (Bellinger et al., 2018). In general, the data availability in GBIF for Collembola from the Neotropics was low. The filtered richness map seemed more accurate, but it still lacks many records of Neanuridae species. Apparently, there are few or no Brazilian Collembola collections in GBIF’s sources. Probably most Neanuridae data are from the Universidad Nacional Autónoma de México. The results of raw and filtered richness and conservation assessments were similar, though filtered data seemed more accurate due to the exclusion of uncertain data.

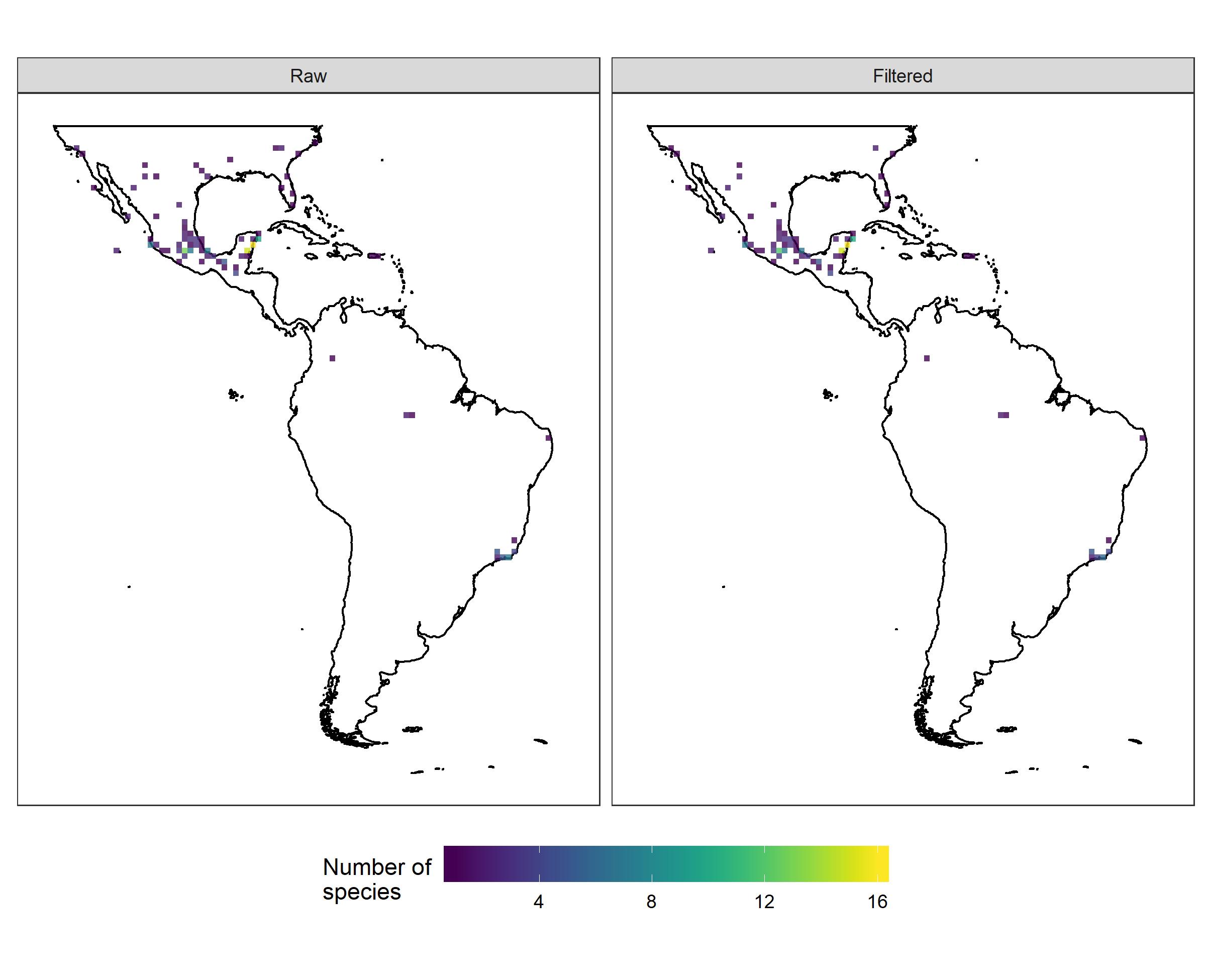

*Oocephalus* (Benth.) Harley & J.F.B.Pastore

*Oocephalus* (Lamiaceae) species previously belonged to two subsections of the genus *Hyptis*, being elevated to generic level by Harley & Pastore (2012). In general, the number of occurrence records available for *Oocephalus* is low, which is likely to be due to this taxonomic change as most of the specimens deposited in herbarium collections are still outdated. The genus, with 20 species (Harley *et al.* 2019), is centered in rocky fields in the Cerrado highlands and Espinhaço range in Brazil (Soares *et al.* 2019). Although both raw and filtered richness maps show similar results, the filtered is more accurate, once records from country centroids were flagged.

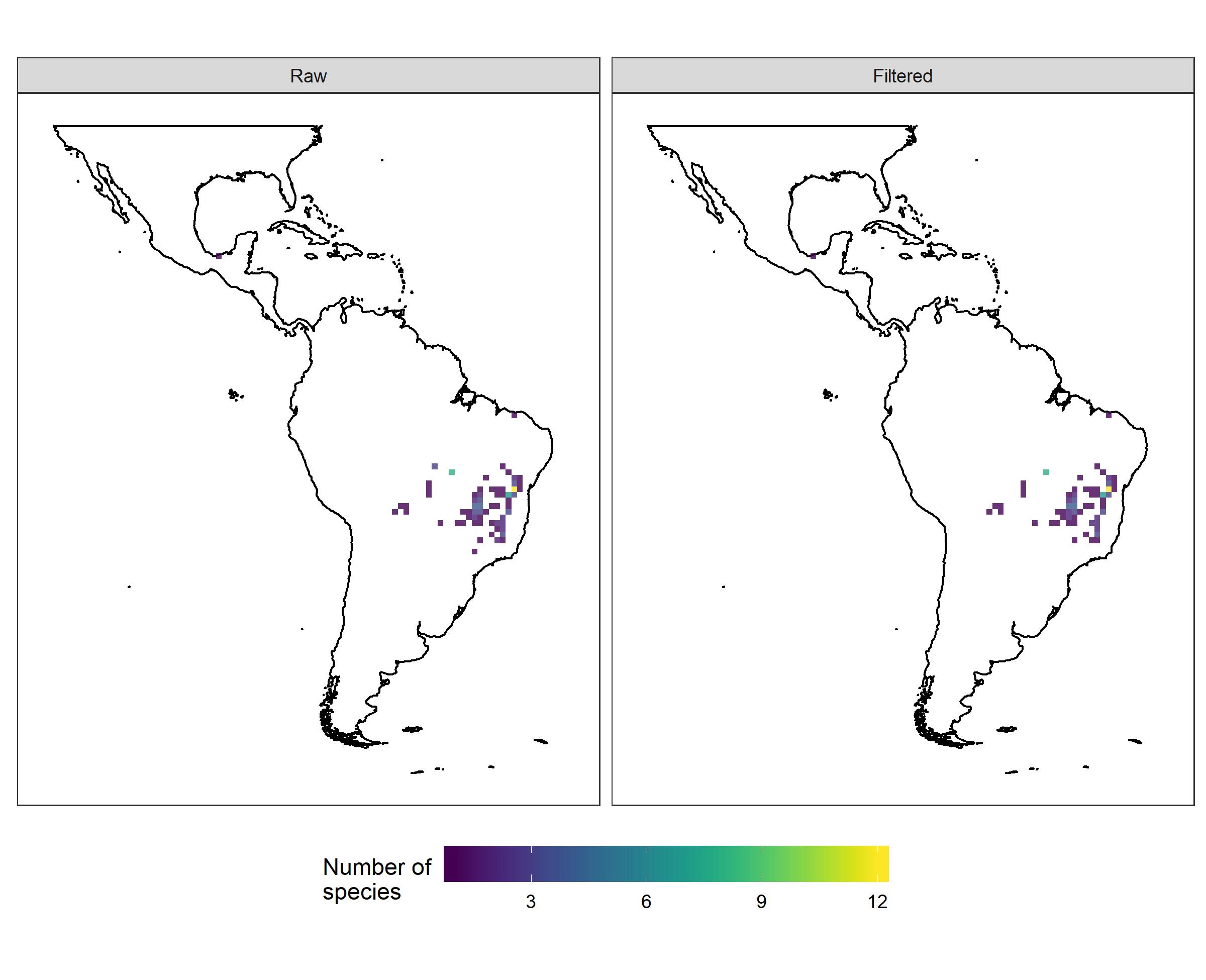

### *Pilosocereus* Byles & G.D.Rowley

*Pilosocereus* is a genus of Cactaceae widely distributed in the Neotropics, with 42 species and 8 subspecies (Hunt *et al.* 2006; Zappi and Taylor 2011). Comparing raw and filtered richness maps, both reflect the known distribution and diversity centers of *Pilosocereus* in eastern Brazil and Mexico (Hunt *et al.* 2006; Calvente *et al.* 2016). However, the filtered map shows more refinement in the distribution of species due to exclusion of outlying records (centroids, duplicates, institutions, sea points). The number of flagged records seems reasonable for the group since “duplicates” increased the total number of records.

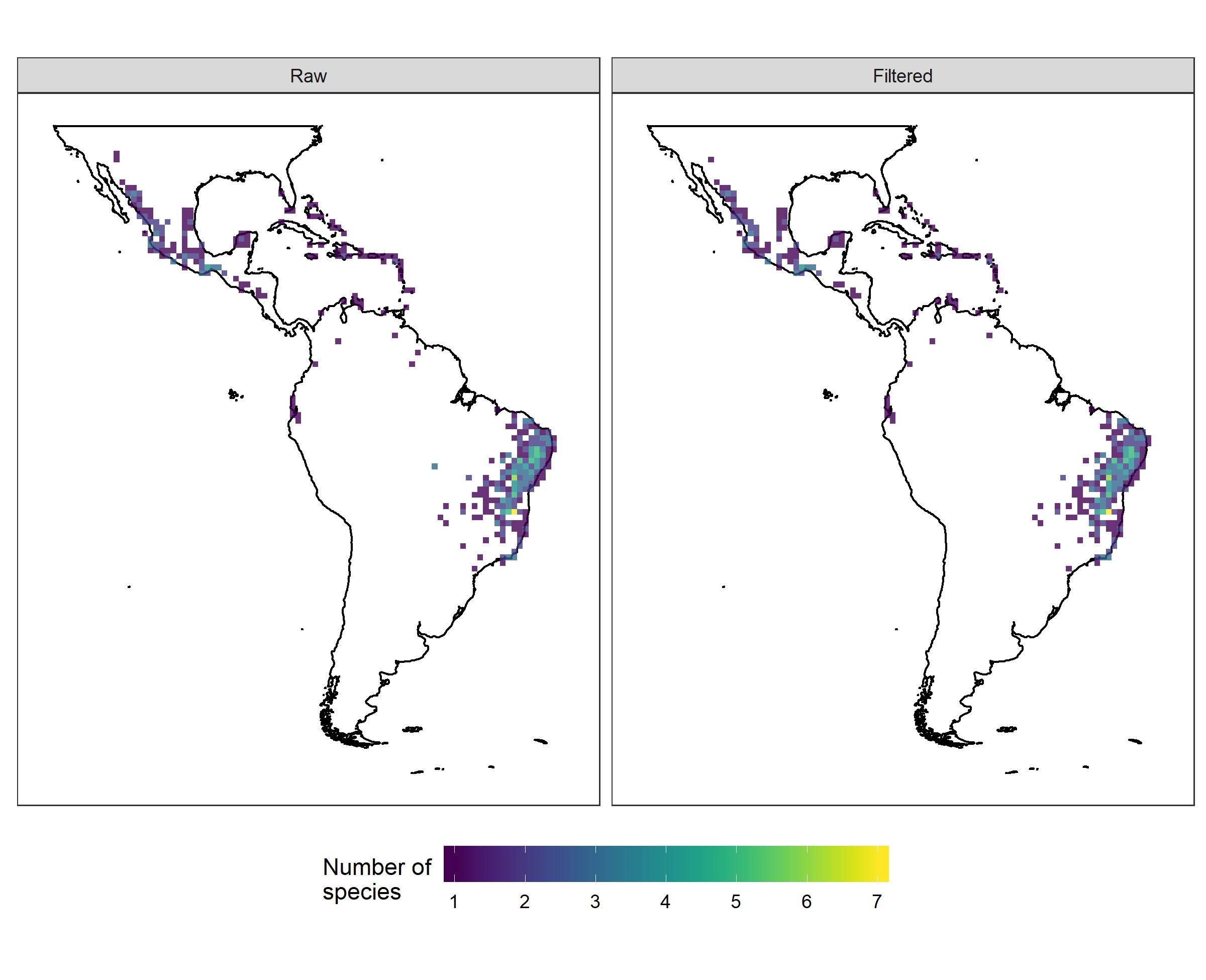

### *Prosthechea* Knowles & Westc.

*Prosthechea* is an epiphytic orchid genus widespread in the Neotropics, comprising approximately 120 species (Govaerts et al. 2020). When comparing raw and filtered richness maps, both reflect the known main diversity centers of *Prosthechea*: Mesoamerica, the Tropical Andes, and eastern Brazil (Govaerts *et al.* 2020, Higgins 2005). Although most of the flagged records were due to duplicates or very old records, a higher refinement on distribution records can be noticed due to the exclusion of some high richness outlier points (e.g. country centroids, capitals, and institutions). These exclusions reduced the alpha diversity range on the map and highlighted the known diversity centers mentioned above, which are areas of mountain formations in the Neotropics. In fact, many species of *Prosthechea* are restricted to high elevation areas, occurring in cloud forests or rocky outcrop vegetation, such as in Mesoamerican mountains, high elevation valleys in the Andes, and the “Serras” in the Brazilian Atlantic Forest. Despite the high refinement of occurrence data achieved with the data cleaning methodologies applied here, taxonomy is still an issue for the genus, due to species complexes, and unclear species delimitation for some taxa.

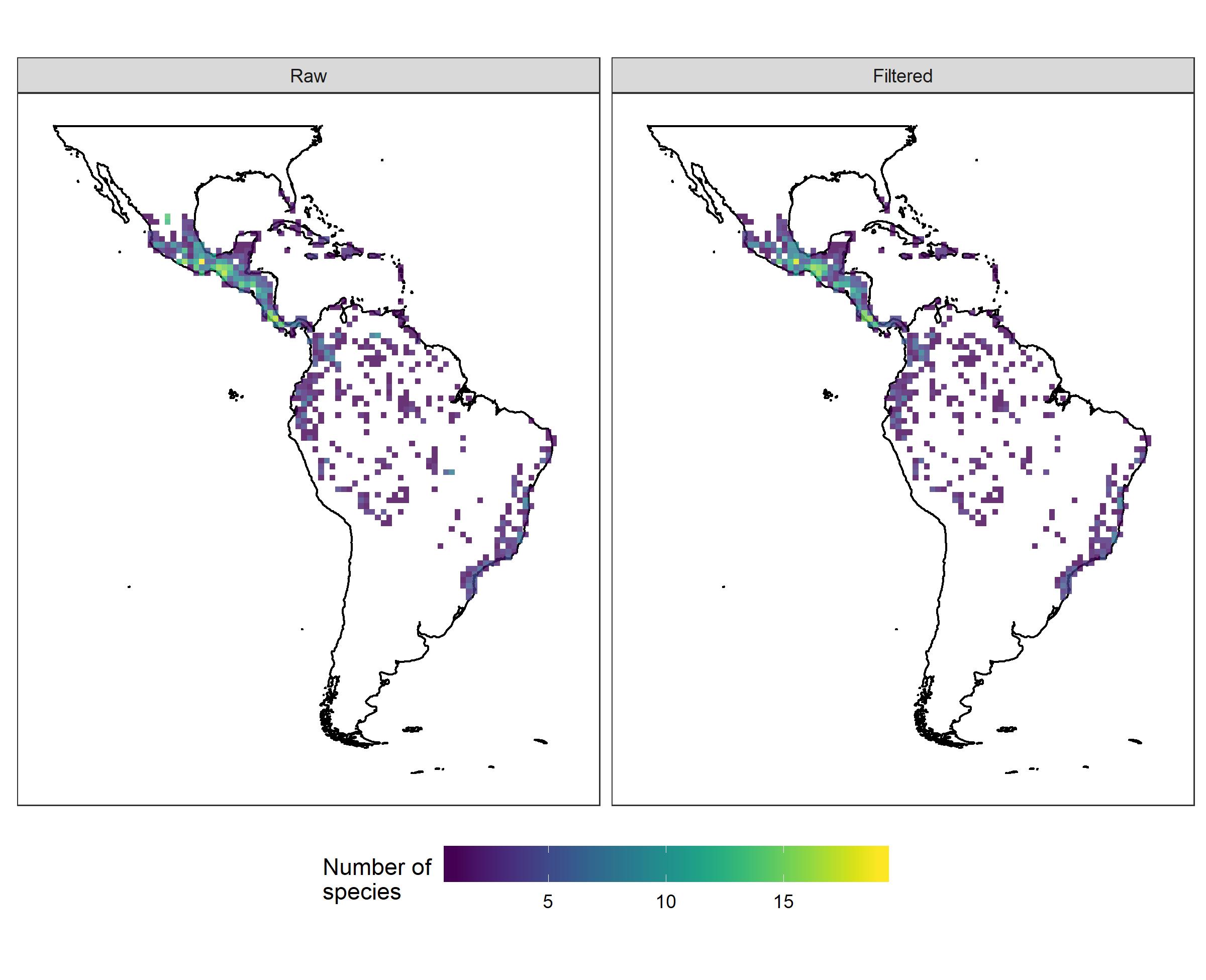

### *Thozetella* Kuntze

*Thozetella* is an asexual Ascomycota genus with c. 22 species worldwide they are saprobes of leaf litter and/or freshwater (Ariyawansa et al. 2015, Monteiro et al. 2016, Perera et al. 2016). There is a concentration of records in the northeast of Brazil (all provided by HUEFS - Herbário da Universidade Estadual de Feira de Santana), and in the south of Mexico (raised on *Inventario taxonómico de los hongos conidiales saprobios del parque Yumka' en Villahermosa, Tabasco* - Heredia-Abarca et al 2018). The genus highlights the gaps of records in GBIF*,* since there are published records from the Brazilian Amazon Forest, Argentina, Venezuela, Cuba and Costa Rica (Silva & Grandi 2013, Monteiro et al. 2016).

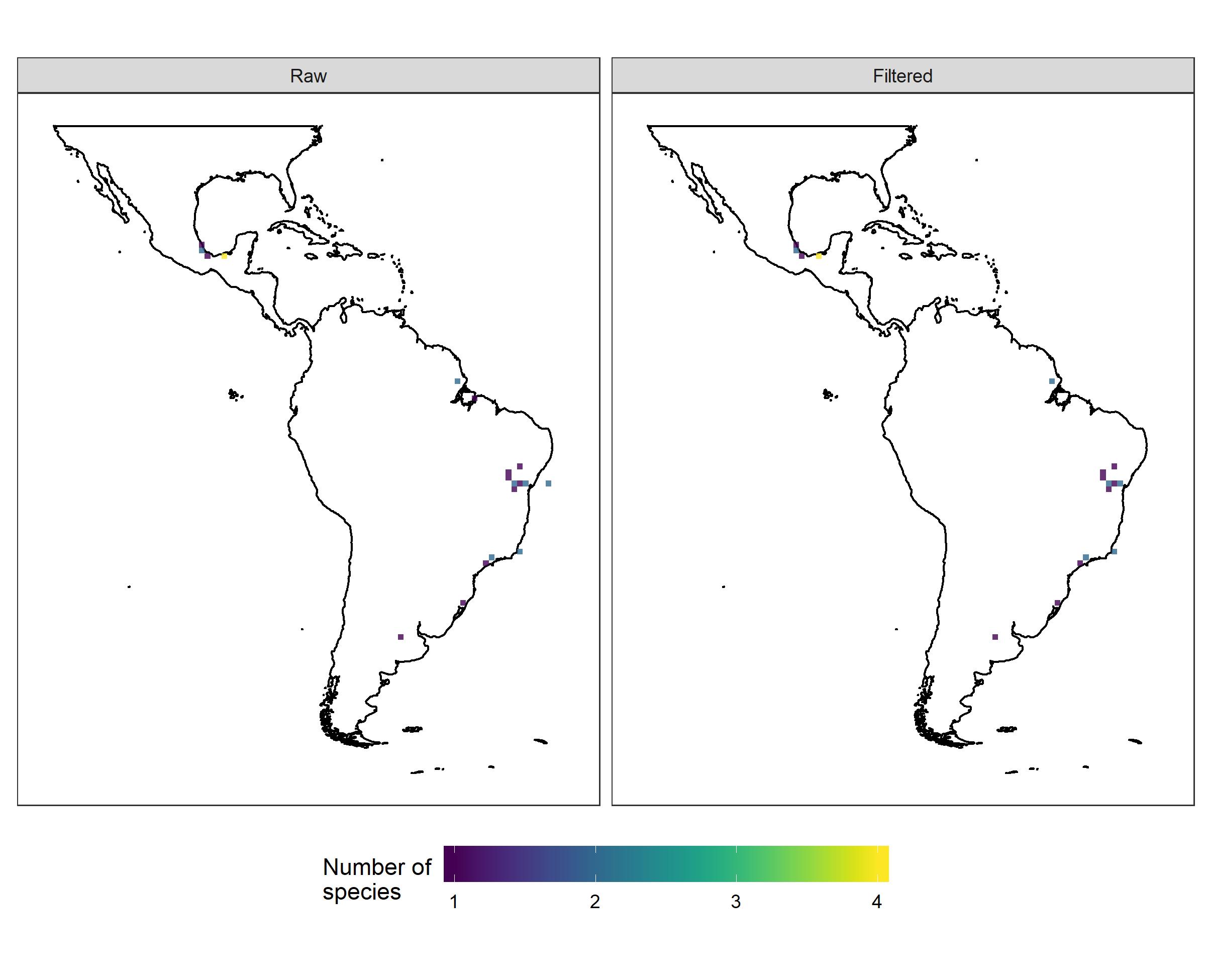

### *Tillandsia* L.

*Tillandsia* is the largest genus in the Bromeliaceae family, with over 700 recognized species, distributed in the Neotropics from the southern United States to Argentina; the highest levels of diversity are found mostly in Central America and in the Andes (Barfuss *et al.* 2016, Gouda & Butcher 2018). The number of flags is acceptable since there were several duplicated records and many located in urban areas, and though several species do naturally occur in urban areas, differentiation from records of cultivated plants would require further analysis. Similarly, many records were not included as they had not been identified to species level or any infra-species categories. Although the cleaned dataset is more precise, the raw data contain more species and may reflect the true situation more closely in terms of species richness.

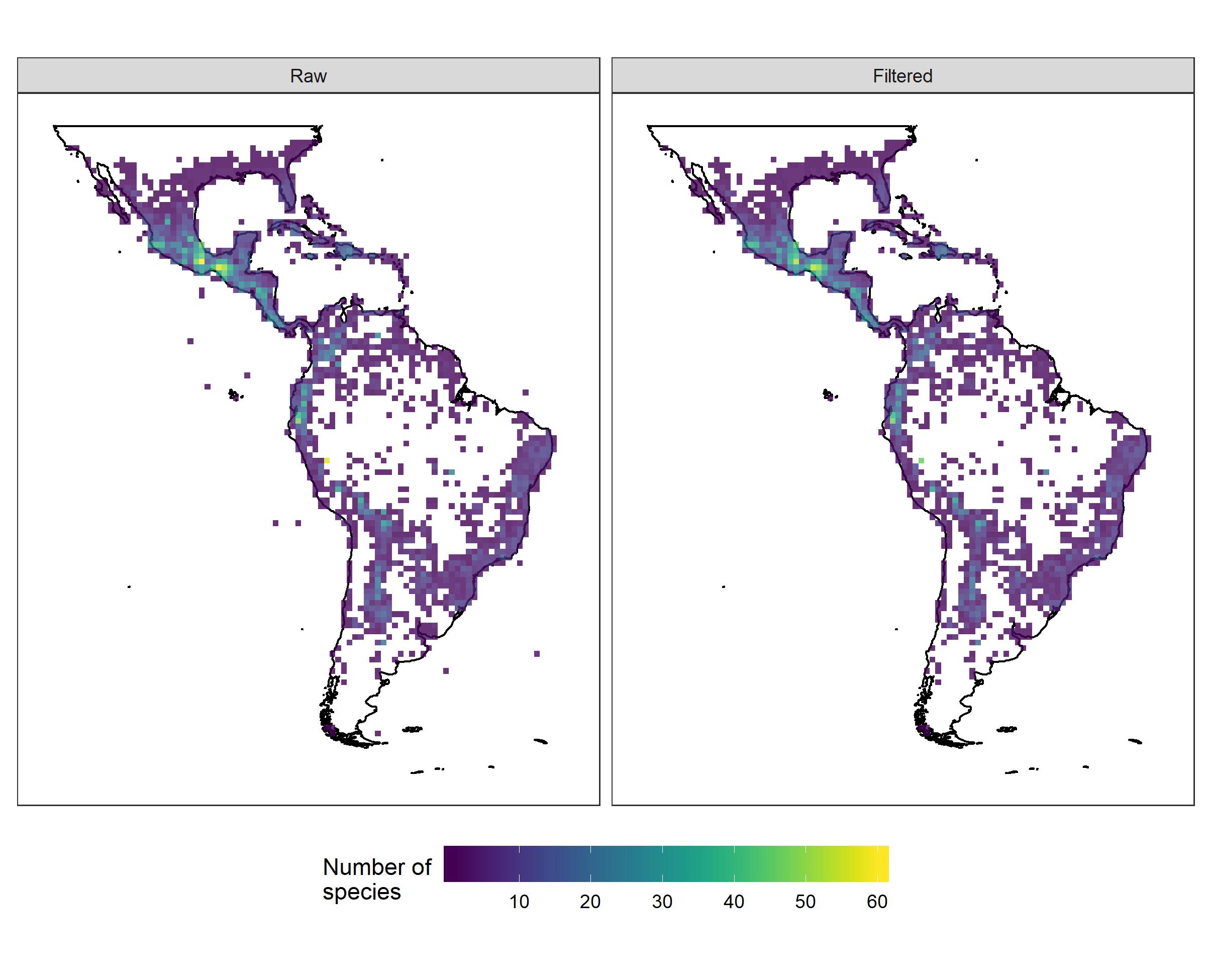

### *Tityus* Koch, 1836

The genus *Tityus* includes the highest number of scorpion species among the family Buthidae, with 220 species described as endemics of the Neotropical region, excluding Chile (Bertani et al. 2008; Lourenço et al. 2015). The richness patterns showed similarities between the raw and filtered data. Even so, the raw dataset appears to be more accurate because it is congruent with the known distribution pattern, with greater diversity of *Tityus* species in Colombia, Ecuador, Venezuela and Central Argentina (Borges et al. 2010, Moreno-Gonzalez et al. 2019). The filtered data can eliminate highest richness regions, as occurred in previous analyzes.

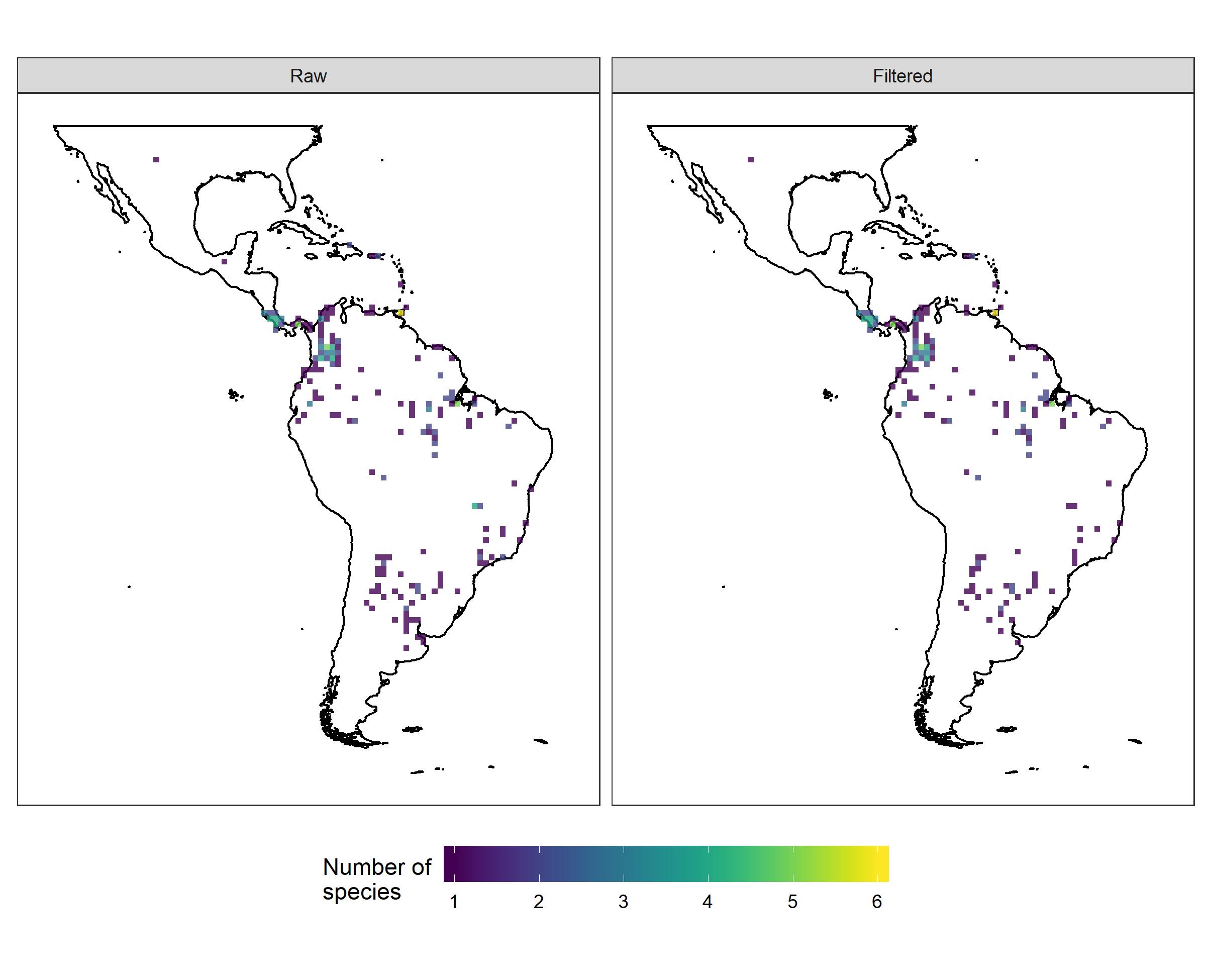

### *Tocoyena Aubl*

*Tocoyena* is a neotropical genus of Rubiaceae, comprising 19 spp. centered in the Cerrado and Atlantic Forest in Brazil (Govaerts *et al.* 2020). Inconsistencies found in the number of species in GBIF raw and filtered data are possibly due to variation of infraspecific taxa identification in herbaria collections. Raw and filtered data showed similar patterns for distribution and conservation assessment, corresponding generally to the known distribution of the genus in the Neotropics (Silberbauer-Gottsberger *et al.* 1992; Borges 2018). However, the filtered map excluded the records for *Tocoyena cubensis*, a species endemic to that region (González-Torres *et al*. 2016; Ulloa-Ulloa *et al*. 2017). The regions highlighted in the maps with higher richness such as northern and northeastern Brazil match the areas previously known as centers of diversity for the genus (Silberbauer-Gottsberger *et al.* 1992).

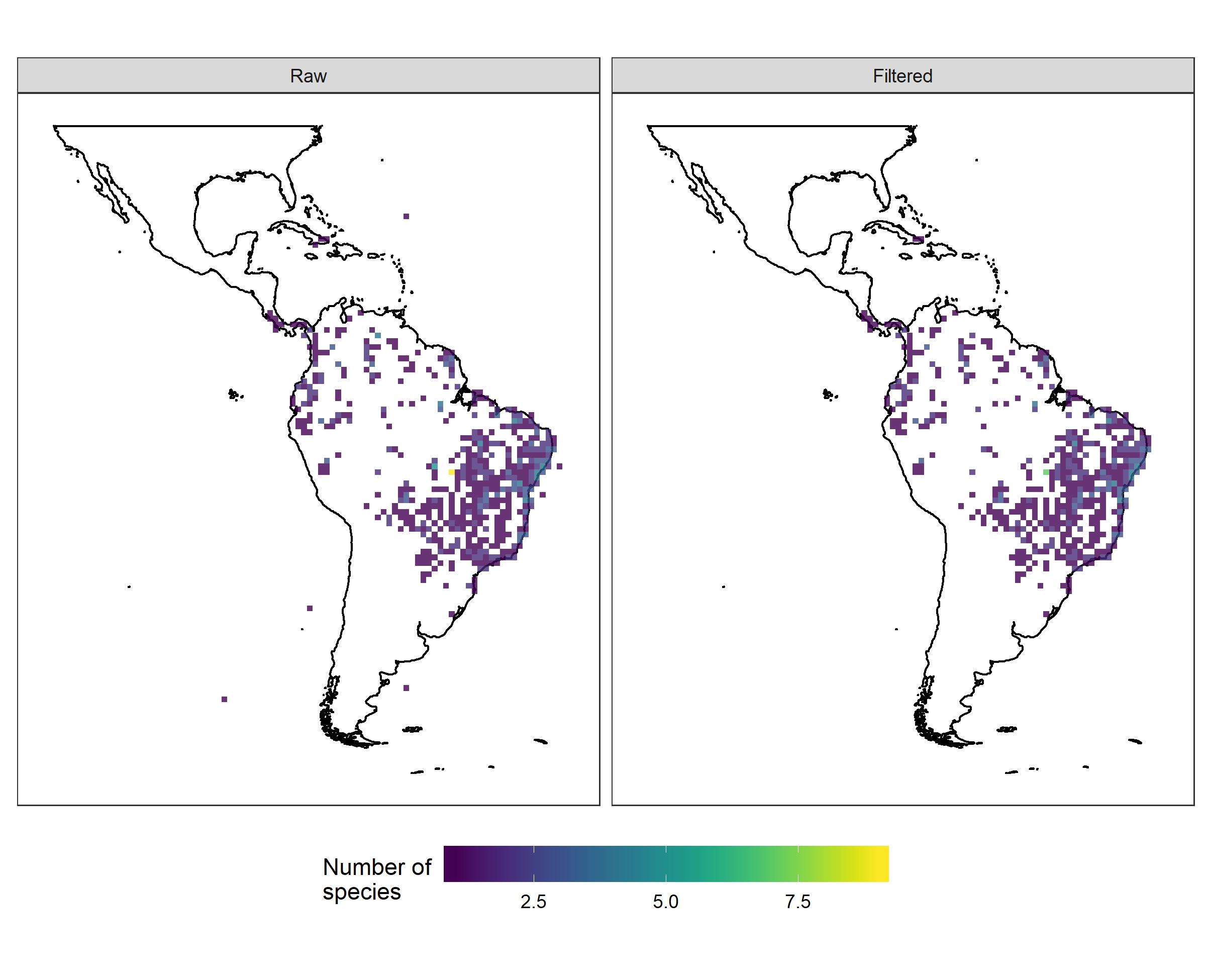
